# Standardising a microbiome pipeline for body fluid identification from complex crime scene stains

**DOI:** 10.1101/2024.08.05.604586

**Authors:** Meghna Swayambhu, Mario Gysi, Cordula Haas, Larissa Schuh, Larissa Walser, Fardin Javanmard, Tamara Flury, Sarah Ahannach, Sarah Lebeer, Eirik Hanssen, Lars Snipen, Nicholas Bokulich, Rolf Kümmerli, Natasha Arora

## Abstract

**Background:** Recent advances in next-generation sequencing have opened up new possibilities for utilizing the human microbiome in various fields, including forensics. Researchers have capitalized on the site-specific microbial communities found in different parts of the body to identify body fluids from biological evidence. Despite promising results, microbiome-based methods have not yet been fully integrated into forensic practice due to the lack of standardized protocols and systematic testing of methods on forensically relevant samples. Our study addresses critical decisions in establishing these protocols, focusing on bioinformatics choices and the use of machine learning to present microbiome results in court for forensically relevant and challenging samples.

**Results:** We propose using Operational Taxonomic Units (OTUs) for read data processing and creating heterogeneous training datasets for training a random forest classifier. Our classifier incorporates six forensically relevant classes: saliva, semen, hand skin, penile skin, urine, and vaginal/menstrual fluid. Across these classes, our classifier achieved a high weighted average F1 score of 0.89. Systematic testing on mixed-source samples and underwear revealed reliable detection of at least one component of the mixture and the identification of vaginal fluid from underwear substrates. Additionally, when investigating the sexually shared microbiome (sexome) of heterosexual couples, our classifier shows promising results for the inference of sexual activity.

**Conclusion:** In our study, we recommend the use of a novel random forest classifier trained on a heterogenous dataset for obtaining predictions from samples mimicking forensic evidence. We also highlight the potential of the sexome for assessing the nature of sexual activities in forensic investigations, while delineating areas that warrant further research. Furthermore, we underscore key considerations when presenting machine learning results for classifying mixed-source samples.

## Introduction

Advances in next generation sequencing technologies have prompted the investigation of the human microbiome for various applications including healthcare, diagnostics and forensics ^1–3^. As more sequencing data is generated from different body fluids and tissues, several key insights have emerged. Among these, bacterial communities have been observed to be body-site specific and bacterial strains to be individual specific ^4,5^. These findings open new avenues for the forensic investigation of biological evidence found at crime scenes, including body fluid identification (BFI) and human identification (ID)^1,6–9^. Body fluid identification from stains provides valuable information to reconstruct criminal events. For instance, in cases of sexual assault, establishing the presence of vaginal fluid or semen can have a significant impact on the investigation.

Recent research has explored the potential of bacterial community sequence data for body fluid identification ^1,6,10,11^. This approach typically targets specific regions of the prokaryotic 16S rRNA gene to describe microbial communities, making it both cost-effective and ideal for forensics, where biological stains are often degraded. Studies have concentrated on forensically relevant body fluids and tissues, including vaginal fluid, semen, and urine, among others. These studies demonstrate that informative, body site-specific markers could be obtained, even when samples were deposited on various substrates or exposed to environmental conditions over an extended period ^1,6,10,11^. Additionally, these studies investigated the presentation of sequencing results as prediction probabilities obtained through various machine learning algorithms like tree-based (*e.g.,* random forests or XG boost) and neural networks among others ^1,6,10^. Despite these encouraging results, microbiome-based analyses are not yet used in routine forensic analysis due to several limitations. These include the lack of standardized laboratory and bioinformatics workflows, knowledge gaps in presenting the analysis output within a probabilistic framework, and the need for further standardization of machine learning output as evidence in court.

The bioinformatics protocols used in previous studies for amplicon sequencing of the 16S rRNA gene are diverse, reflecting the wealth of approaches and software available for microbiome analyses. However, this variation in turn highlights the need for standardised workflows in applied settings like forensic casework ^12,13^. Amongst the different options available, two important bioinformatic decisions concern the clustering of read data (or not) and whether to combine publicly available data from different 16S rRNA gene regions in order to obtain larger reference datasets for comparative analyses and machine learning. Given the inherent biases associated with each 16S rRNA gene region, it is unclear whether heterogeneous sequence combination introduces new biases ^14–16^.

Once these bioinformatic decisions are addressed, the second aspect for forensic application is the inclusion of microbiome-based predictions as evidence. Forensic analyses typically employ a probabilistic framework for interpretation by the court, allowing the findings to be evaluated within the context of their evidentiary value^1,17^. Despite encouraging results from previous studies, machine learning tools are yet to be integrated into the forensic routine. This is due to gaps such as limited body fluid categories in the training sets and limited testing on forensically relevant samples. Biological evidence from crime scenes is often found in degraded conditions, deposited on various substrates and present as mixtures of two or more body fluids, among other possibilities^18^. While previous studies have tested machine learning classifiers on aged samples and samples on substrates, these samples were typically high input, unlike forensic evidence. Moreover, the performance of these classifiers on mixtures remain understudied ^6,9,10^.

As a first step in our study, we begin with addressing the bioinformatics aspect. We compared the resolution of OTUs (operational taxonomic units) clustered at 97% identity threshold versus ASVs (amplicon sequencing variants) for BFI purposes. Next, using OTUs generated with closed reference clustering, we combined data from nine different datasets, including public and novel datasets. These datasets were generated by sequencing different regions of the 16S rRNA gene (V1V3, V3V4, V4 and V4V5 regions). Subsequently, a random forest classifier was trained on this combined dataset comprising six training classes, namely, saliva, semen, skin from hand, penile skin, urine and vaginal/menstrual fluid. Our training dataset included forensically relevant body fluids/tissues from both commonly studied body fluids as well as understudied ones, providing a comprehensive representation for the classifier’s training.

As a last step in study, we tested our random forest classifier on mock forensic samples such as mixed-source samples and samples from substrates with two levels of complexity. The first level included mock samples generated in controlled conditions in the laboratory and the second level included mock samples representing forensic evidence. Such evidence in sexual assault cases generally includes clothing items such as underwear from victims and swabs from the urogenital areas of victims and suspects^19–22^. We included underwear samples from women and vaginal, semen and penile skin swabs from couples for investigating the sexually shared microbiota (sexome). We collected extensive metadata for both these datasets including details on sexual activities and personal hygiene. In summary, we propose the use of OTUs from different 16S rRNA gene regions, producing a heterogeneous machine learning training dataset that is more robust to cross-study analyses. In addition we demonstrate the advantages of systematically testing our classifier for inferring body fluids/tissues from samples mimicking forensic casework.

## Results

### Body site clustering patterns from OTUs and ASVs are highly similar

We first compared the clustering patterns between closed reference OTUs (97%) and ASVs for samples from saliva, semen, skin from hands, menstrual blood and vaginal fluid. We used a V4V5 dataset of 42 samples containing 3965 ASVs and 2647 OTUs among the five body fluid/tissues studied. We generated PCoA plots using weighted Unifrac (UF) (Figure 1, a and b) and Bray Curtis (BC) dissimilarity distances (supplementary material S1, FigureS1.1).

**Figure 1.**
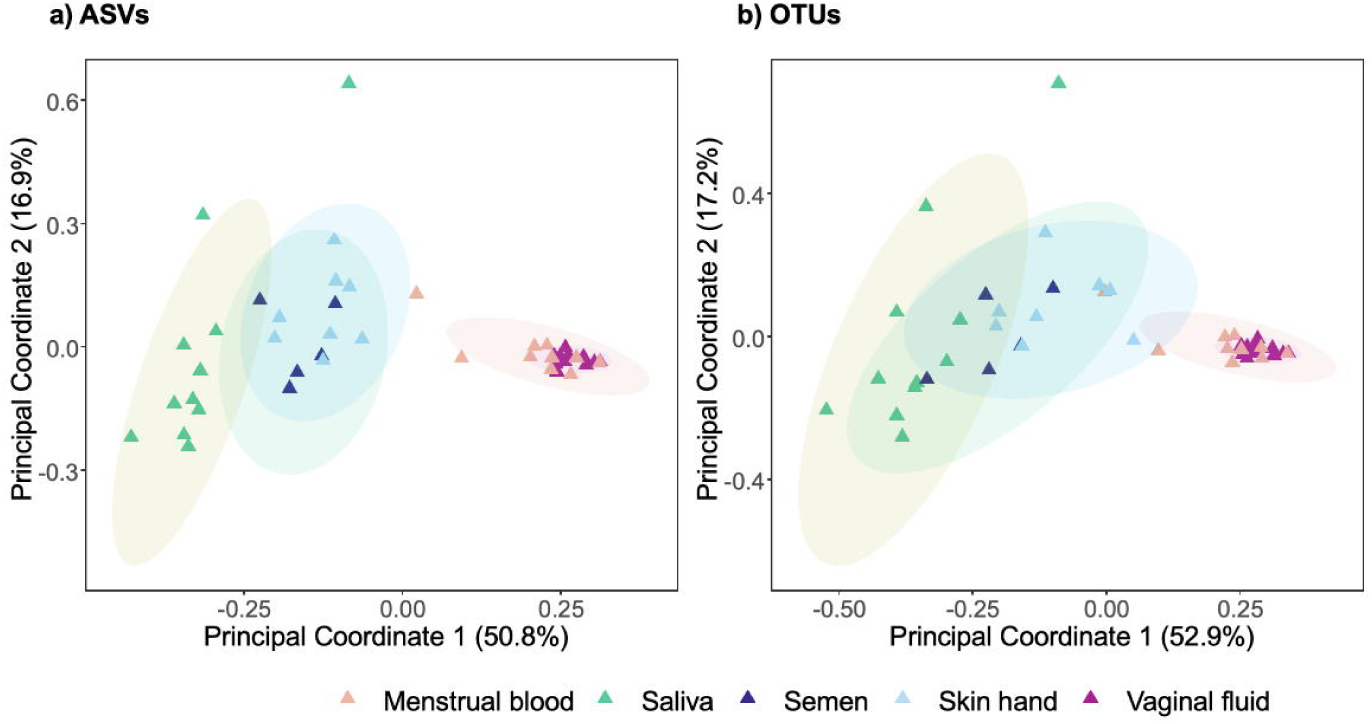
Principal coordinate analysis plots (based on the weighted unifrac distances) showing clustering of body fluid/tissue samples. **a)** ASV data for samples from Dobay et al. (n=42, PERMANOVA F_4,41_ = 16.15, r^2^ = 0.61, p = 0.001), b) OTU data clustered at 97% for samples from Dobay et al. (PERMANOVA F_4,41_ = 15.95, r^2^ = 0.61, p = 0.001). Body fluid/tissues are colour coded.

We observed that the samples clustered according to body site, for both OTUs and ASV data. In all PCoA plots, irrespective of distance metric, significant effects of body fluid/tissues on clustering patterns were observed (ASV PERMANOVA weighted UF, F_4,41_ = 16.15, r^2^ = 0.61, p = 0.001 and OTU PERMANOVA weighted UF, F_4,41_ = 15.95, r^2^ = 0.61, p = 0.001, ASV PERMANOVA BC F_4,41_ = 6.15, r^2^ = 0.38, p = 0.001 and OTU PERMANOVA F_4,41_ = 7.64, r^2^ = 0.43, p = 0.001). Pairwise comparisons for Figure 1 revealed three main body fluid/tissue clusters were observed: (i) saliva, (ii) semen together with skin from hand, and (iii) vaginal fluid with menstrual blood samples (supplementary material S2). The first two principal coordinate axes in the OTU analysis explained a slightly higher proportion of variance (70.1%) compared to the principal coordinate axes in the ASV analysis (67.7%). Regression analyses on PC1 and PC2 coordinates for both distance metrics demonstrated strong correlations between the ASV and OTU coordinates for both axes (PC1, r^2^ = 0.979 and PC2, r^2^ = 0.987)(supplementary material S1, Figure S1.2). Overall, we found similar clustering patterns for ASVs and OTUs suggesting that both can be utilized for body fluid identification.

### Samples from different 16S rRNA gene regions cluster similarly

Next, we assessed whether OTUs for different 16S rRNA gene regions provided comparable distinction across body fluids/tissues for body fluid identification. OTU clustering was conducted using a closed reference clustering method, a robust and computationally faster approach compared to other options like fragment insertion. We used a subset of 48 samples from the Zurich Institute of Forensic Medicine dataset (herein referred to as Zurich dataset) from 6 different body fluids/tissues. The 48 samples comprised sequence data from V1V3, V3V4, and V4V5 regions (16 samples for each region) using similar protocols detailed in supplementary material S3. We observed highly similar clustering patterns using weighted Unifrac irrespective of the gene region analysed (Figure 2a-c). Regression analyses between the coordinates of PC1 and PC2 of different gene regions again revealed strong correlations (r^2^- values between 0.88-0.98, supplementary material S1.3), confirming the highly congruent clustering. Given the highly similar clustering patterns, we agglomerated the OTU abundance tables from all 48 samples into one PCoA plot (Figure 2d). With the combined data set, the body fluid/tissue clustering pattern remained congruent with those in Figures 2a-c. Crucially, body fluid/tissue was a significant driver of clustering patterns (PERMANOVA, p-values < 0.05, see Figure 2 legend for exact p-values), while the 16S rRNA gene region was not (PERMANOVA F_2,45_ = 0.33, r^2^ = 0.01, p = 0.972). In summary, our analyses showed that data from different 16S rRNA gene regions can be combined without affecting the clustering of body fluids/tissues.

**Figure 2.**
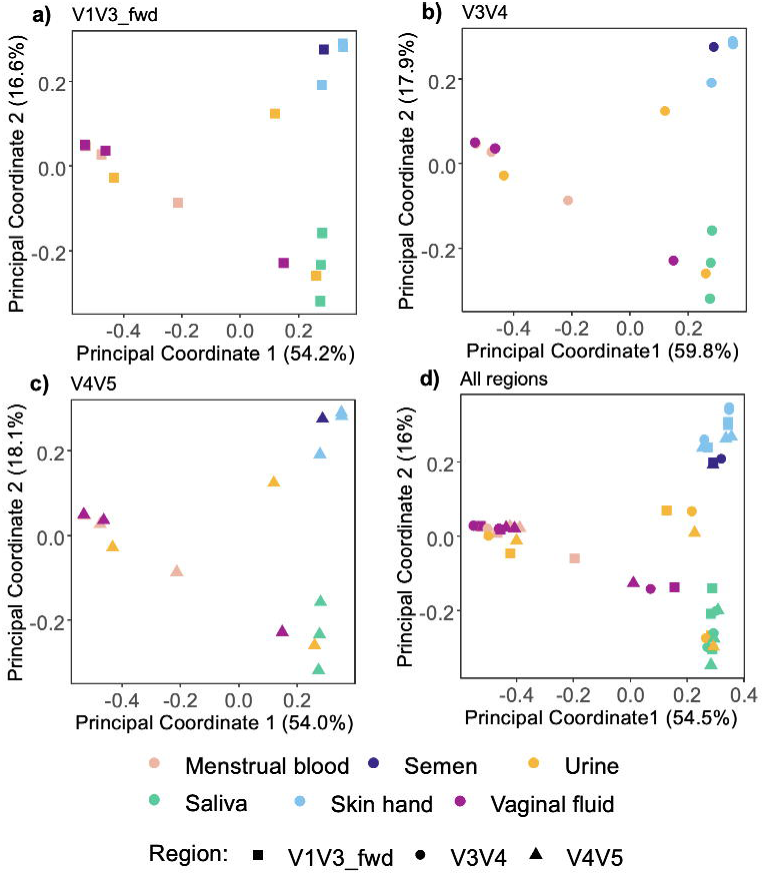
Principal coordinate analysis plots for data from the different 16S rRNA gene regions using OTUs (97%) and weighted UniFrac distances. **a)** V1V3 data (n=16, PERMANOVA F_5,10_ = 3.57, r^2^ = 0.64, p = 0.003), **b)** V3V4 data (n=16, PERMANOVA F_5,10_ = 4.89, r^2^ = 0.71, p = 0.002), **c)** V4V5 data (n=16, PERMANOVA F_5,10_ = 4.76, r^2^ = 0.7, p = 0.001), and **d)** All three regions combined, V1V3, V3V4 and V4V5 (n=48, PERMANOVA F_5,42_ = 15.07, r^2^ = 0.64, p = 0.001). Body fluid/tissues are colour coded and the 16S rRNA gene regions are depicted by different shapes.

### Random Forest Classifier trained on a heterogenous dataset achieves high prediction accuracies

In order to generate a heterogeneous training dataset, we combined read data from 9 different studies conducted in different laboratories with diverse protocols targeting one or more regions of the 16S rRNA gene (V1-V3, V3-V4, V4 and V4-V5)^9,11,22–24^. The reads originated from 457 samples from six body fluid/tissues namely, saliva, semen, skin from hands, penile skin, urine and vaginal swab. In total, there were 6455 OTUs across all fluids/tissues. As expected, significant effects of body site were observed (F_6,450_ = 55.33, r^2^ = 0.42, p = 0.001, supplementary material S1, Figure S1.4). In addition, the 16S rRNA region was also found to be significantly associated with microbiome composition, albeit with lower effect size(F_4,452_ = 6.82, r^2^ = 0.06, p value = 0.001 for both variables, supplementary material S1, Figure S1.4).

In the next step, we trained a random forest classifier on the OTU compositional data of six classes namely, saliva (n = 99), semen (n = 57), skin from hands (n = 80), penile skin (n = 20), urine (n = 42) and vaginal/menstrual fluid (n = 159). The training was done using an 80-20 train-test split resulting in a training dataset of 365 samples and a testing dataset of 92 samples. A total of 6455 OTUs were obtained across all samples, however, the most discriminatory OTUs were extracted using the recursive feature elimination. We found that 281 out of 6455 OTUs reached overall accuracies > 80% (supplementary material S1, Figure S1.5). These OTUs were then used for model training. Subsequently, we applied the trained model on the 92 test samples and the reliability of the classifications were assessed using F1 scores as the performance metric. Both weighted average F1 scores and F1 scores per class were analysed. A high weighted average F1 score of 0.89 was obtained across the 6 training classes (saliva, semen, skin from hand, penile skin, urine and vaginal swab). The F1 scores per class varied whereby higher F1 scores were observed for saliva, semen, skin from hands, penile skin and vaginal swabs (0.85-1) than for semen and urine (0.70 and 0.46 respectively) (Figure 3a). Although F1 scores provide a good overview on classifier performance, assessing individual prediction probabilities for each sample and all training classes is critical in applications like forensics. In most studies conducted so far, accuracy of a classifier is assessed based on the highest prediction probability, whatever the value is and without setting a threshold. With this approach, 88% of the 92 test samples were correctly predicted and 12% were categorised as misclassified. As illustrated in Figure 3b, among the misclassifications, three semen samples were incorrectly predicted as vaginal, with varying probabilities ranging from 0.33 to 0.89. The remaining misclassifications mostly occurred amongst fluids originating from the same or proximally located sites like semen, penile skin and urine, which is an expected outcome given the potential for cross-contamination via microbial transfer.

**Figure 3.**
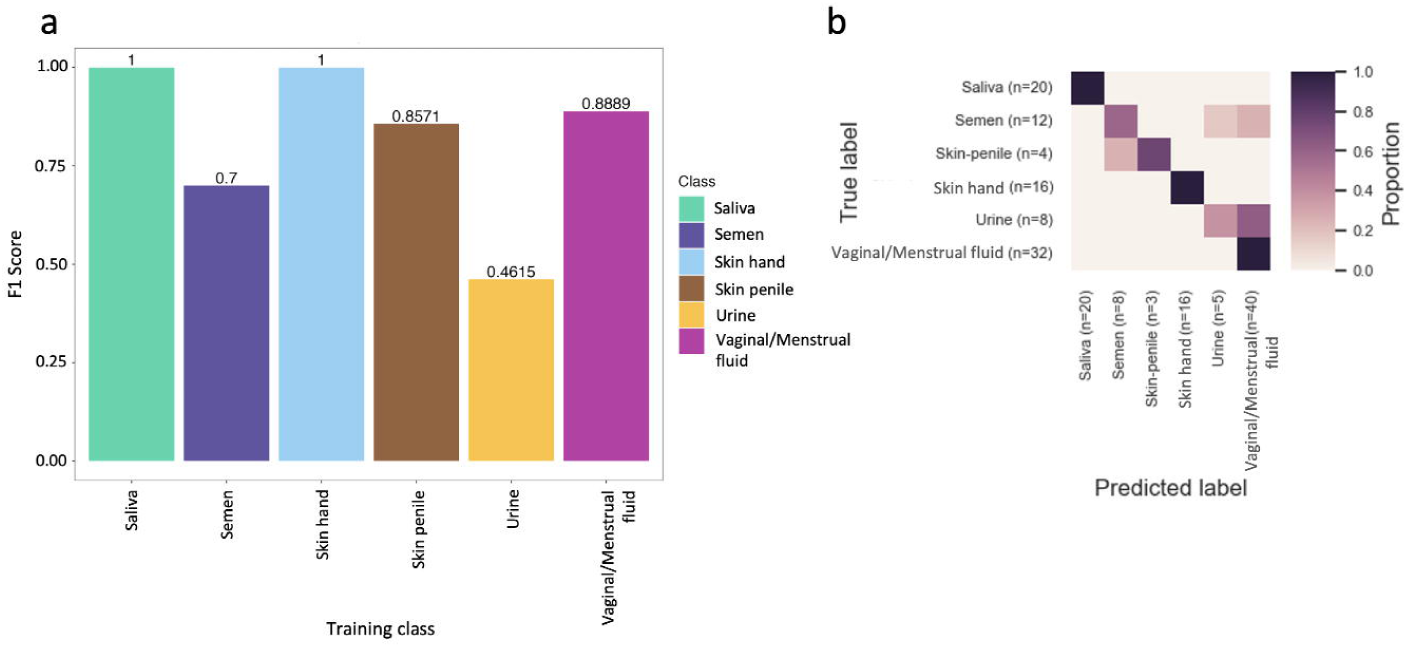
Classifier performance of the classifier trained on 365 samples on 92 test samples. **a)** Barplots depicting F1 scores per class **b)** Confusion matrix for the 92 samples with sample sizes per class, the exact prediction probabilities are reported in supplementary material S4.

Despite these overall encouraging results, some samples exhibited low probability values for the predicted class (below 0.5) or displayed similar probabilities for more than one training class. For instance, sample *A007,* which was correctly predicted as semen with a probability value of 0.38, exhibited a similarly low probability value 0.34 for urine (data in supplementary material S3). As semen and urine pass through the urethral tract in males, it is possible that the similar values reflect a mixed composition of these fluids. Nonetheless, such probability distributions could pose challenges in forensic casework.

As an alternative approach to basing prediction solely on the class with the highest probability, we tested inferring body fluids/tissues based on probability thresholds. If the maximum probability fell below the threshold, the sample was left unclassified. Given the high microbial biomass observed in saliva, skin from hand, vaginal swab and penile skin classes and based on the distribution of prediction probabilities, a threshold of 0.7 was chosen. A lower threshold of 0.5 was selected for fluids/tissues known to exhibit lower microbial biomass such as semen and urine. The application of thresholds is a more conservative approach as it reduces both the proportion of correctly classified (from 88% to 77%) and misclassified (from 12% to 6%) samples. Simultaneously, it introduced the new category for unclassified samples (17%), which enabled us to acknowledge the uncertainty in predictions especially for low bacterial biomass samples and samples originating from proximal body sites like e.g. semen, penile skin or urine. In summary, a high number of correct predictions were obtained in most cases with or without thresholding and whenever un- or misclassifications occurred they typically concerned proximally located sites.

### High prediction probabilities obtained for blind single-source samples and controlled mixtures

Once we were satisfied with the classifier performance on the test set, we used the same parameters to train an extended classifier using all 457 samples and an independent (blind) dataset was used for assessing classifier performance. A total of 337 OTUs from the total of 6455 OTUs were selected for model training using recursive feature elimination (supplementary material S1.6). To test this new classifier, we used blind controls from the Zurich dataset, consisting of 47 single-source samples comprising 10 samples from saliva, 9 samples from skin from hand/forearm, 8 samples from semen and 20 vaginal/menstrual fluid samples. The random forest classifier performed extremely well, yielding high F1-scores for all categories (0.85-1.00) (supplementary material S1.7). The probabilities for each sample in the blind dataset was visualized using boxplots (Figure 4a). Overall, 95% and 82% of the 47 samples were correctly classified without and with thresholds respectively (supplementary material S5). With thresholds, saliva, skin from hands and vaginal swabs samples were generally correctly classified (80-100%), however, only 37% of the semen samples were classified correctly. As observed previously, unclassified samples predominantly comprised those with low microbial biomass and from proximal sites. Despite the unclassified samples, no samples were misclassified showing that the classifier exhibited reliable performance upon assessment with an independent dataset.

**Figure 4.**
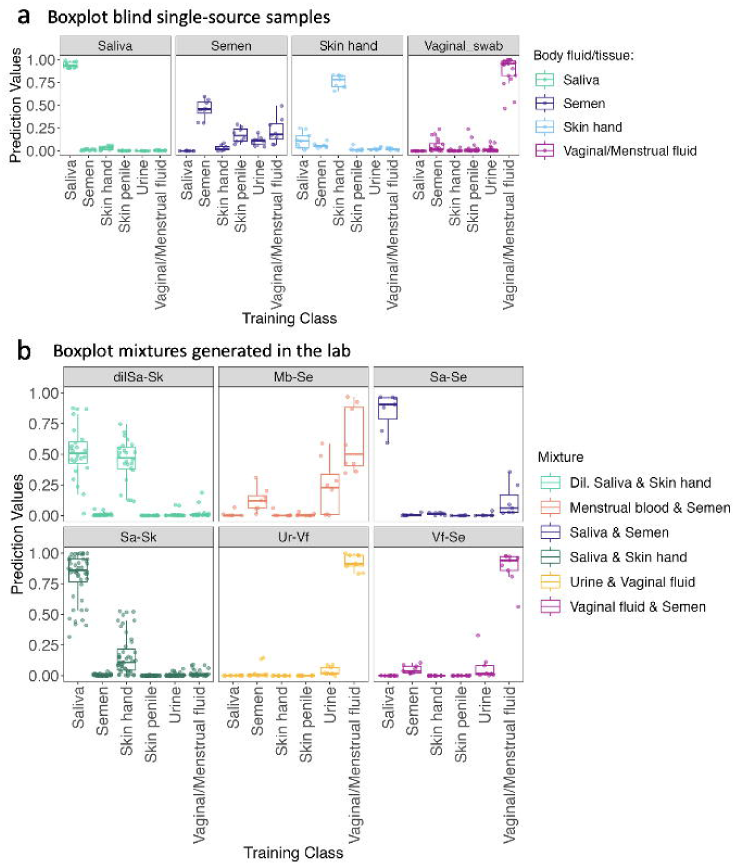
Boxplots depicting prediction probabilities of blind single-source control and mixed source samples generated in the laboratory. **a)** Blind single-source samples faceted according to the body site. **b)** Mixtures generated in the lab faceted according to mixture composition. Each data point refers to a sample and each sample has a probability value per class. Dil. Saliva & Skin refers to 1:10 diluted saliva deposited on skin.

Next, we tested the classifier for its ability to identify body fluids from mixed-source samples since such samples are commonly encountered in forensic casework. This set comprised 104 mixed-source samples generated in the lab by mixing 2 sources/body fluids and for these samples, the distribution of probabilities was explored (Figure 4b and supplementary material S6). The outcomes for this analyses were distinguished as follows; (i) where both the mixture constituents display the top two highest probabilities, (ii) where one of the mixture constituents displays the highest probability and (iii) where neither of the two mixture constituents display the highest probability. We observed that in 81.7% of the 104 samples, both constituents had the highest prediction probabilities. These cases comprised a large number of the saliva and skin samples (71 out of 104 samples). These results are expected from saliva and skin mixture samples as the two constituents are comparable in microbial load and biomass unlike other mixtures such as menstrual blood and urine. Among the remaining mixtures, and with the exception of two samples (16.3% of the 104 samples), at least one of the components had the highest prediction probability. Since these samples mainly comprised samples where there was a discrepancy in the microbial load between the two constituents (menstrual blood-semen, saliva-semen, urine-vaginal fluid, and vaginal fluid-semen), the component that was generally identified was vaginal/menstrual fluid or saliva. Lastly, in 1.9% samples, neither of the two mixture constituents displayed the highest prediction probability. These samples consisted of menstrual-semen mixtures and exhibited the highest probability for the class urine. To determine whether there were significant differences among the means of the different classes, an ANOVA with Tukey post-hoc analysis was conducted corroborating the interpretations of our findings (supplementary material S7). In summary, we observed that in most cases, the classifier could reliably identify at least one of the mixture constituents, especially the ones that are high in microbial load.

### The classifier can detect expected traces in underwear and sexome samples accurately

In order to assess the suitability of the classifier in forensic casework, the classifier was tested on non-controlled mock samples, mimicking the samples collected during forensic casework. The predicted class was determined to be the body site with the highest prediction probability.

The first set of non-controlled mock samples consisted of underwear samples collected from 10 women after wearing the garment for 24h. Prediction probabilities for all 10 underwear samples were visualized as boxplots (Figure 5a). All samples were predicted as vaginal swabs and the prediction probabilities ranged between 0.43-0.89. We observed some signatures for semen and expected minor signatures for skin from hands (supplementary material S8). Therefore, the classifier could detect microbial taxa for the vaginal body site from underwear as the substrate.

**Figure 5.**
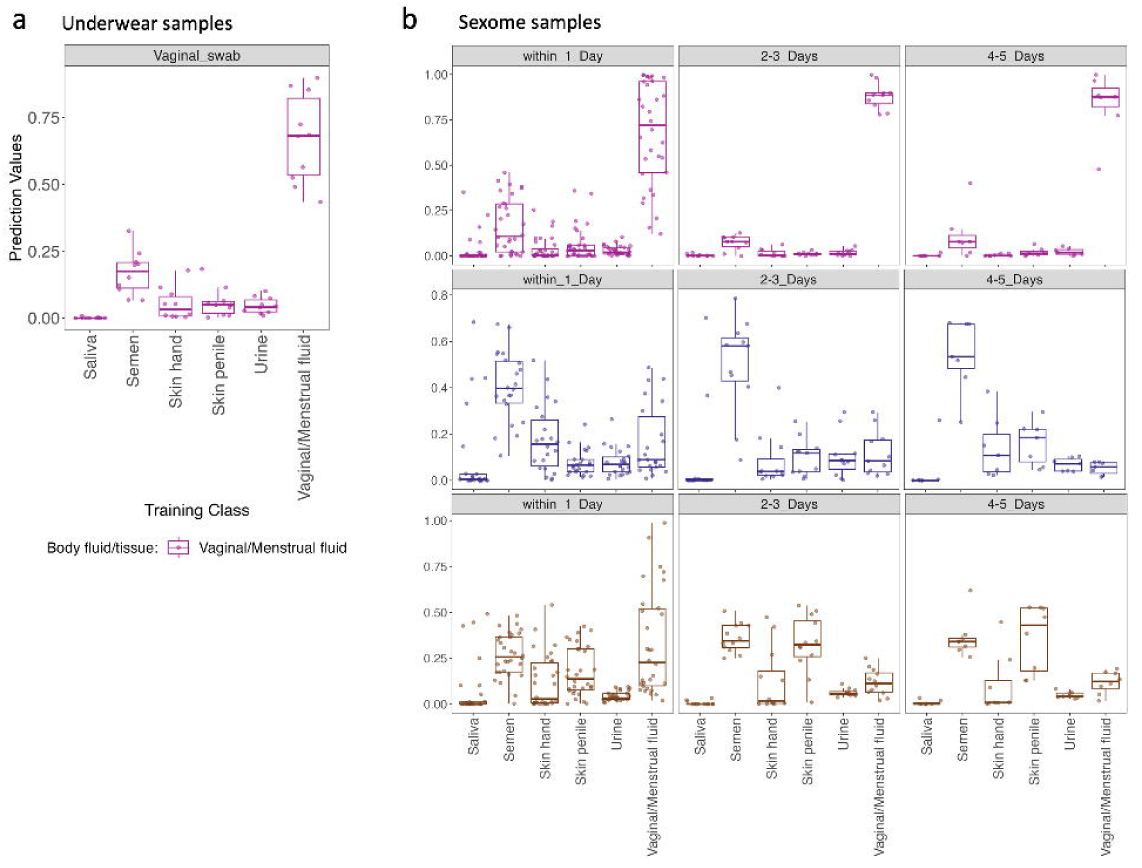
Boxplots displaying prediction probabilities for inferring urogenital microbiota from underwear and sexome samples. **a)** Underwear samples. **b)** Sexome samples based on days elapsed between sexual activity and sample collection. Each data point refers to a sample and each sample has a probability value per class.

The second set of non-controlled mock samples were obtained as part of the Zurich study of urogenital samples from 22 heterosexual couples. The goal of the study was to investigate sexually shared microbiota (sexome) and to establish whether sexual activity can be inferred through the detection of mixtures of body fluids/tissues. Such a possibility would be useful for the analyses of sexual assault cases, where detecting both male components, such as penile skin or semen, and female components from the vaginal area on collected evidence would be beneficial. The dataset consisted of a total of 139 samples collected/deposited on swabs including vaginal fluid, semen and penile skin. For all these samples, metadata on sexual activities and personal hygiene was also collected. Across all days, and without applying any thresholds for the predictions, we observed 89% of vaginal fluid samples, 67% of semen samples and 28% of penile skin swabs predicted as the respective class. (supplementary material S8). Subsequently, we further investigated the prediction probabilities after categorising the data into three groups based on the days since sexual activity: 1 day, 2-3 days and 4-5 days. As illustrated in Figure 5b, prediction probabilities indicated mixing of the male and female components especially in samples collected within 1 day after sexual activity. However, these probabilities become less indicative of such mixing as more days pass since sexual activity. Nonetheless, in one vaginal sample (V2t1) semen traces and in a penile skin sample (P16t2) vaginal traces could be detected until 4 days after sexual intercourse. In addition, the prediction probabilities observed are also indicative of the nature of sexual interactions. For samples where oral traces were expected (as specified in the metadata), saliva could be detected up to 4 days after sexual activity. As an example, in the vaginal swab, semen and penile skin samples from couple 21, microbial signatures indicative of saliva in addition to skin and vaginal fluid could be detected (Figure 6). Interestingly, the prediction probabilities indicative of oral or sexual intercourse (for instance saliva or vaginal fluid on penile skin) were lower on the fourth day after sexual intercourse. In summary, the ability to detect the intermixing of traces and nature of sexual interactions could provide important insights in forensic cases.

**Figure 6.**
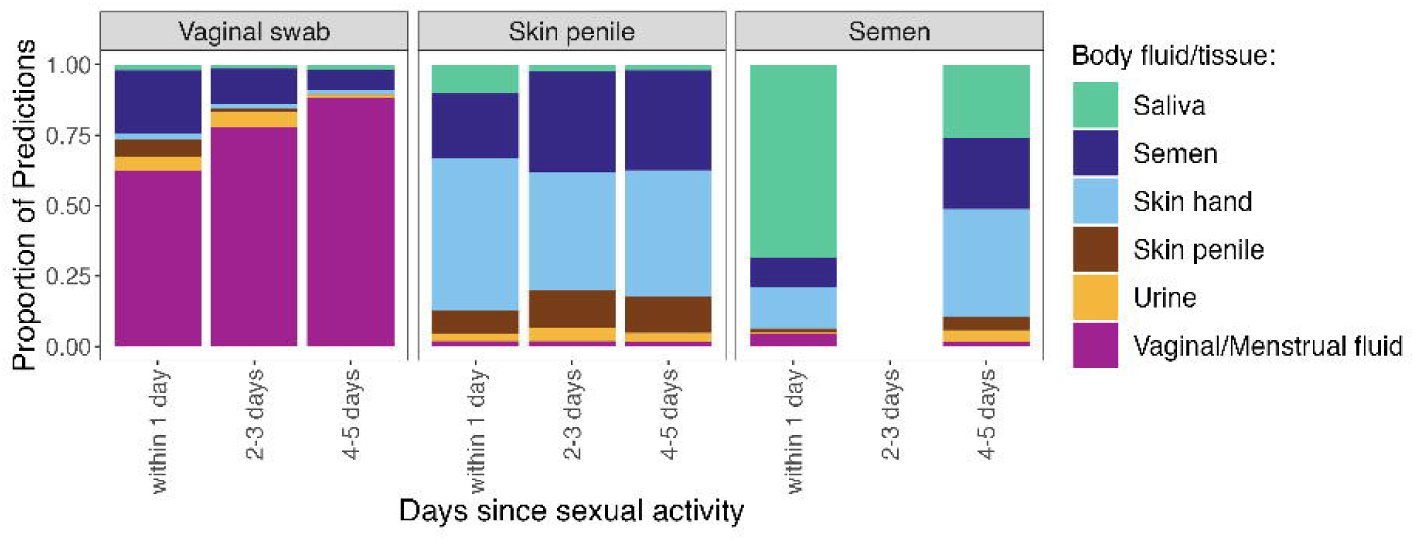
Sexually shared microbiota from couple 21 where oral, semen and penile traces were indicated. The semen sample for the 2-3 days was excluded during quality filtering as it had less than 1000 reads.

### Analysis of informative taxa provides explanations for the predictions

Finally, we evaluated the informative taxa using feature importance scores that can be extracted from RFCs to identify and rank the features that contributed most to the model predictions. Top 10 informative taxa per training class were selected and a total of 45 OTUs out of the 337 OTUs were selected and visualized (Figure 7). As demonstrated in Figure 7, the most informative taxa used to classify saliva, skin from hand, penile skin and vaginal and menstrual fluid consist of distinct and characteristic OTUs, explaining the high prediction probabilities obtained for the correct classes in our study. For instance, amongst the taxa with highest feature scores in saliva we find *Veillonella* and *Fusobacterium* whereas skin from hand was characterized by OTUs from *Cutibacterium*, *Pseudomonas*, *Staphylococcus*, *Corynebacterium* and *Enhydrobacter*. In the case of urine, among the top 10 informative taxa we find two OTUs, *Pelomonas* and *Enterobacter*, that are characteristic of this body site as well as other OTUs also found at other body sites. Overlapping OTUs that hinder the distinction of body sites are generally found for body fluids/tissues that are proximally located such as urine (from females) and vaginal/menstrual fluid. Nonetheless, some characteristic OTUs like *Enterobacter* and *Prevotella* were found in urine. In addition, the differences in relative abundance of *Lactobacillus, Gardenerella and Streptococcus* led to a distinction in vaginal swabs and urine. Similarly, the distinction of penile skin and semen was challenging as well since the most informative markers for each of these body fluids/tissues overlap, reflecting the proximal location of these two sites. However, the difference in frequencies of some shared OTUs such as *Prevotella*, *Porphorymonas* and *Corynebacterium* among others resulted in the overall distinction between these two classes. Some overlapping taxa were present with similar frequencies between classes, providing explanations for our misclassifications and probability distributions between semen and penile skin, urine and semen from males and urine from females and vaginal/menstrual fluid. For instance, vaginal/menstrual fluid samples were enriched with *Lactobacillus*, *Streptococcus* and *Gardenerella*, however, these OTUs were also observed in low frequencies in semen or urine samples. Similarly, overlapping *Lactobacillus* and *Pelomonas* OTUs were seen between semen and urine classes. Therefore, the investigation of the top informative taxa provides possible explanations for the findings in our study.

**Figure 7.**
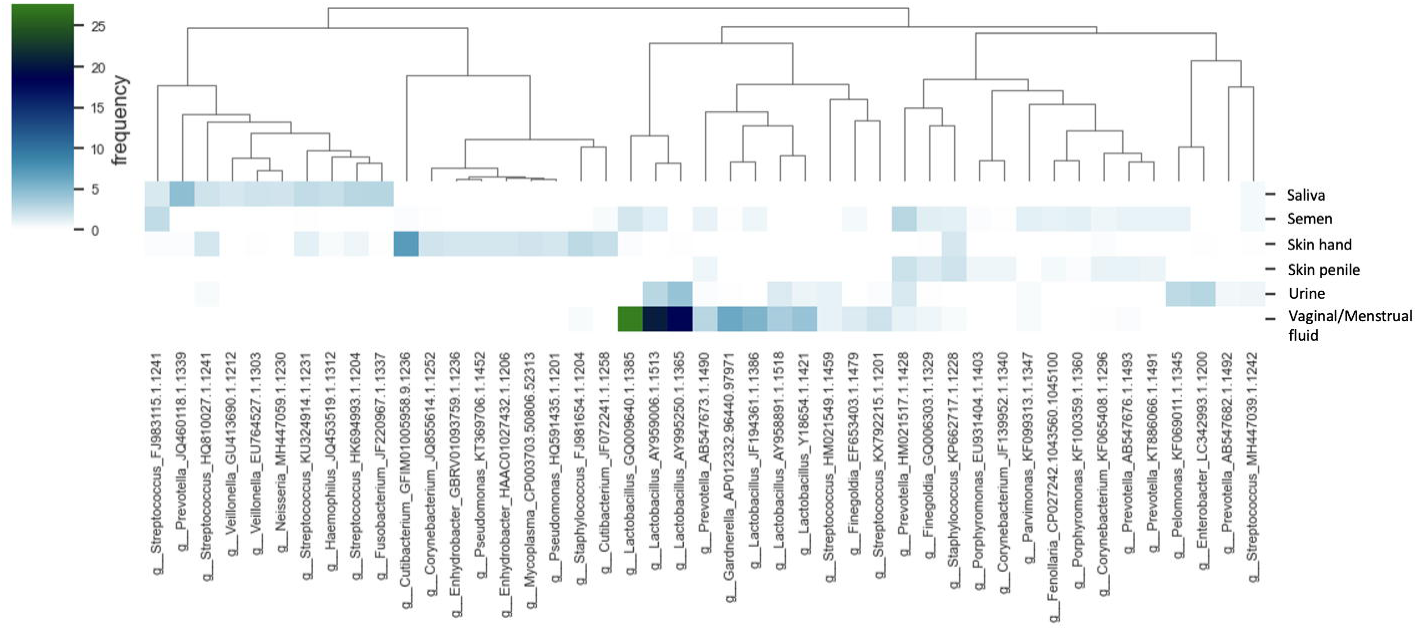
Heatmap of 10 most informative OTUs for each training class with genus level taxonomy. 45 out of the 337 OTUs are displayed.

## Discussion

In our study, we addressed two key aspects for standardisation of protocols for forensic utility, namely, bioinformatics and inclusion of microbiome-based predictions as evidence for court. Regarding the bioinformatics aspect, previous comparisons of OTUs and ASVs have demonstrated that ASVs may enable a more precise characterisation of microbial diversity, as well as improved detection of rare taxa^3,15,25–28^. At the same time, ASVs have been shown to cause errors in relative abundance calculations by providing inflated numbers of ASVs and genome splitting^29^. Overall, it appears that the advantages and disadvantages of ASVs and OTUs depend on the goal. The results in our study highlight that both operational taxonomic units (OTUs) and amplicon sequence variants (ASVs) from targeted 16S rRNA gene regions offer comparable resolution for forensic body fluid identification (BFI). In studies focusing on goals like biomarker discovery or detection of (rare) bacterial strains, even deeper resolution obtained through long read or shotgun sequencing may be needed ^8,30,31^. However, the comprehensive identification of diversity as well as rare taxa is not needed for forensic body fluid identification. In this study, we demonstrate that while ASVs allow for the distinction of body sites, the comparison of studies based on ASVs is more challenging. Importantly, OTU clustering opens avenues to harness the combined power for different 16S rRNA gene regions.

Our analyses on the integration of data from different 16S rRNA gene regions showed that sample clustering patterns were driven primarily by the body site rather than the region. These results are congruent with those of other studies investigating the potential of combining data from different 16S rRNA gene regions ^15,16^. The authors in Jones et al.(2022) demonstrated that combining data from different 16S rRNA gene regions offers a higher taxonomic resolution and a more comprehensive view of bacterial diversity compared to using single regions susceptible to biases^15^. Our classifier trained on the combined dataset therefore effectively identifies informative taxa for body fluid classification across the four regions incorporated here (V1V3, V3V4, V4 and V4V5). Moreover, Tackmann et al. (2018) and colleagues showed that machine learning (ML) classifiers trained on large combined datasets displayed higher accuracies than classifiers trained on smaller datasets comprising a single region of the 16S rRNA gene^16^. Therefore, such a comprehensive dataset is not only beneficial in the forensic context, but also in other applications seeking to characterise the human microbiome and predicting disease states^32^.

Using machine learning algorithms advances us to the next forensic aspect addressed in this study: the inclusion of microbiome-based predictions as evidence for court. This aspect encompasses obtaining reliable prediction probabilities from forensically relevant samples and presentation of these probabilities in an evaluative report. As the first step to obtain prediction probabilities, we trained a Random Forest Classifier (RFC). Our evaluation of F1 scores per class for assessing the reliability of our classifications revealed that our findings were comparable or slightly higher than those reported in previous studies ^1,16^. The slight improvement in our F1-score for semen in comparison to Wohlfahrt et al. (2023) could be due to the heterogenous training dataset employed here, combining various 16S rRNA gene regions instead of only the V4 region alone. Another potential explanation could be the use of slightly varied training classes in both studies. While our study lacked the extensive sample sizes of Tackmann et al. (2018), our F1 scores were highly comparable for the overlapping body fluids/tissues, namely, saliva, skin from hand and vaginal^16^. In addition, we included other understudied forensically relevant samples like semen, penile skin and urine, albeit with smaller sample sizes. We observed variations in per class F1-scores which could be due to technical factors like sample sizes or biological factors such as microbial diversity, biomass or physiological sample location (Figure 3a). These factors may also explain the lower prediction probabilities observed for urogenital samples when compared to saliva, skin and vaginal/menstrual fluid (Figure 3b). Analysis of the blind single-source samples using the classifier trained on the entire dataset of 457 samples yielded overall higher F1-scores compared to the test set (supplementary material S1.7). The increased per class F1-scores could potentially indicate the significant impact of the larger sample sizes on classifier performance.

In evaluating misclassifications to assess the reliability of our classifications, we found that our classifier encountered difficulties in accurately classifying certain body fluids, particularly semen samples. Potential explanations for these challenges include: low microbial diversity and biomass, and contamination from proximate sites. Another important explanation for the vaginal swab false positives in some semen samples is the absence of metadata regarding the sexual activities of these donors. Thus, it remains unclear whether these samples were genuinely single-source at the time of collection and samples with the above limitations will rarely achieve high F1-scores and accuracy.

Even though F1-scores provide an insightful overview of classifier performance, our findings highlight the importance of evaluating prediction probabilities for each class and sample. An avenue to increase confidence in our correct predictions is to employ thresholds for classifying single-source samples and introduce an “unclassified” category. This approach serves to minimize both false positives and false negatives, a critical consideration in applied settings ^33,34^. While Diez-Lopez et al. (2019) previously adopted a constant arbitrary threshold of 0.7 across all body fluids/tissues, we adopted flexible thresholds due to varied microbial biomass of our training classes^10^. A threshold of 0.5 was used for semen and urine samples and a threshold of 0.7 was applied to other single-source samples. For the three semen misclassifications from the test set, three semen samples were unclassified and two misclassified out of the five samples when thresholds were applied. Interestingly, upon applying thresholds, the same sample sequenced with the V4V5 region fell into the unclassified category whereas the V3V4 sample remained misclassified. One possible explanation could be the lower number of semen samples from the V3V4 region compared to the number of semen samples from the V4V5 region in the training dataset.

Although thresholding showed promise in single-source samples, such an approach is not possible with mixed source samples. Our findings from the controlled mock mixtures generated in the lab showed that disentangling both constituents of all mixtures was challenging and prompted further considerations. A worthwhile strategy is to implement thresholds for the cumulative probabilities associated with the relevant body fluids and tissues (Figure 6). However, a forensic scientist must not be privy to information on relevant fluids/tissues during evaluation. Therefore, when establishing a standardized workflow, it may be beneficial to include a predefined set of relevant body fluids for specific case types, such as sexual assault cases. Such an approach can then be consistently applied across all sexual assault cases, irrespective of the specific details of any particular case, ensuring a robust and uniform framework. In addition, our analyses also highlight the importance of setting a lower threshold to determine “undetected” body fluids/tissues among the training classes. Given these complexities, other advanced tools like multi-class classifiers or Bayesian Networks offer a promising avenue for forensic use. Bayesian Networks have been previously used for forensic DNA STR analysis and could potentially consolidate all probability distributions obtained for a sample and provide a resultant likelihood ratio ^35^.

Our analyses from the underwear samples and sexome samples showed that the classifier can detect signatures from substrate samples and that samples from couples could indicate the time frame and nature of sexual activity. Underwear has been previously studied in the context of vaginal health, however, such samples could potentially offer another avenue for detecting sexual activities ^36,37^. However, our study was not specifically designed to explore this aspect and further investigations on the underwear microbiome are necessary to assess its forensic applicability. The findings from the sexome samples yielded further interesting insights for forensic utility. Our findings were consistent with a case report focused on health related aspects of the microbiome where microbial taxa indicative of sexual activity could be detected in vaginal and penile skin samples for up to 4 days^38^. In addition, our investigation uncovered the presence of oral traces also detectable up to 4 days after sexual intercourse, offering additional understanding of the nature of sexual activities. Knowledge of these time frames coupled with analyzing the proportional decrease in male and female components therefore provides potential for establishing timelines of sexual activities in case work. Our results demonstrate that the sexome can provide additional lines of evidence or strengthen the evidentiary value of other traces as suggested by our findings and those of Dixon et al. (2023)^39^. Despite their limited sample size of 6 couples in total, the authors demonstrated unique male and female bacterial signatures before intercourse and significant transfer of the female microbiome to male partners after unprotected sex. Therefore, further research including larger cohorts and pre-coital and post-coital samples from the same couples is essential to fully leverage the sexome’s potential in forensic casework. Most of the current sexome studies primarily focus on investigating vaginal health and dysbiosis ^2,40,41^, the findings from health and forensic research offer valuable knowledge that benefits each other. In a recent study by Carter et al. (2024), the strain-level identification of bacterial taxa shared between couples is shown in a health context^31^. Such identification can also be applied to forensics for donor identification. Notably, our study represents, to the best of our knowledge, the first application of machine learning to predict sexome samples of this scale. Our findings show that generation of larger sample cohorts studying the transmission, persistence and recovery of the microbial signatures between sexually involved individuals is warranted. Results from such cohorts may benefit both forensic science and health-related research such as transmission pathways of urogenital bacteria ^31^.

## Conclusion

In our study, we highlight important decisions on two key aspects, bioinformatics and evaluative reporting, to integrate microbiome-based analyses in forensic casework. Moreover, we propose a novel RFC trained on a heterogenous dataset including forensically crucial and underexplored body fluids/tissues like semen, urine and penile skin and systematically test it on complex samples. Our study, therefore, provides an outlook on employing machine learning in forensics along with investigating the sexome in future studies. We provide resolution on some aspects and highlight imminent gaps that should be addressed for microbiome analyses to be incorporated in forensic body fluid identification.

## Methods

### Description of all datasets, controls and mock samples

In this study, we combined data from 9 different microbiome datasets comprising 4 unpublished studies and 5 published studies targeting the V1-V3, V3-V4, V4 and V4-V5 16S rRNA gene regions. Across all studies, a total of 788 samples from 7 different body fluids/tissues namely saliva, semen, menstrual blood, vaginal fluid, urine, and skin from palm/forearm and penile skin were included. Figure S1.8 in the supplementary material S1 describes the number of samples in all categories.

### Data from published studies

Read data for the V3V4 region, V4V5 region and V4 region from single source control and mixed source samples were obtained from the studies of Hanssen et al. (2017), Dobay et al. (2019) and Leeber et al. (2023) respectively ^11,25,26^. Further, V4 region read data from the studies of Williams et al. (2019) and Meisel et al. (2016) were downloaded using q2-fondue in QIIME2 (v2023.5)^27,28^. The sample ids of all samples will be provided upon request.

### Data generation for unpublished studies: Zurich datasets

The four unpublished datasets were generated in our lab at the Zurich Institute of Forensic Medicine and are referred to as Zurich or Zurich datasets throughout the manuscript and the metadata. These datasets targeted the V1V3, V3V4 and V4V5 regions of the 16S rRNA gene. One V1V3 dataset (Zurich dataset 1) was generated using slightly different protocols and the remaining V1V3 samples, V3V4 and V4V5 samples (Zurich dataset 2) were generated using the same protocols. All four datasets consisted of different numbers of control, mixed-source and substrate samples.

### Sample collection for Zurich datasets

Participants contributing to the Zurich datasets were recruited during the course of the different studies conducted between 2020-2023. All participants were provided with an informed consent form. All single source samples were collected directly on cotton swabs or pipetted onto swabs from collection tubes/containers (see details in the supplementary material S3). Controlled mock mixture samples were prepared in the lab. In the non-controlled mock samples, underwear samples were collected after women had worn the garment for 24 h and sexually shared microbiome (sexome) samples were collected and deposited on swabs. Appropriate blanks and negative controls were included. Donors for the underwear and sexome samples were asked to fill out questionnaires including details on sexual activities and personal hygiene.

### DNA extraction and quantification for Zurich datasets

DNA extraction for all samples was conducted using one of the following kits with minor modifications in the protocols: QIAamp BiOstic Bacteremia DNA Kit (Qiagen, Germany), DNeasy PowerSoil Pro Kit (Qiagen, Venlo, Netherlands), DNA purification from buccal swabs (Spin protocol) from the QIAamp DNA Mini Kit (Qiagen, Venlo, Netherlands) and PrepFiler BTA Automated Forensic DNA Extraction Kit (Thermo Fisher Scientific, Waltham, MA, USA). The swab was either directly added to the Powerbead tubes or the cotton was separated from the wood with a sterile scalpel and placed into a 2 ml tube (Investigator Lyse & Spin Basket Kit, Qiagen, Venlo, Netherlands). The samples were lysed according to each protocol and the lysate was separated from the swab by centrifugation. The extraction was then carried out following the manufacturer’s protocols with minor modifications.

DNA quantification was performed using one of the following protocols: Quantus Fluorometer (Promega, Inc., Madison WI, USA) or a SYBR green assay with the FemtoTM Bacterial DNA Quantification Kit (Zymo Research, Irvine, CA, USA) or the protocol described by Seashols-Williams et al. (2018) using the PerfeCTa SYBR Green SuperMix (Quantabio, Beverly, MA, USA) and the ZymoBIOMICS™ Microbial Community DNA Standard D6306 (Zymo Research, Irvine, CA, USA). Cycle numbers for the amplicon PCRs were decided based on the Ct numbers obtained during the qPCR.

### Library preparation and sequencing Zurich datasets

Bacterial communities were obtained by amplifying V1V3, V3V4 and V4-V5 regions of the 16S rRNA gene. The V1V3 region was amplified using modified F27/R534 primers with spacers, the V3V4 region using F341/R806 and the V4V5 region using modified F806/R926 primers. The exact primer sequences with spacers are provided in the supplementary material S3. All primers were ordered from Sigma (Darmstadt, Germany) or Microsynth AG (Balgach, Switzerland). The PCR protocol and cycling conditions were optimized per region and Zurich dataset and are provided in the supplementary material S3. PCR products were cleaned-up using the AMPure XP beads (AMPure XP, Beckman Coulter). Single-size selection was performed for the V1-V3 and V3-V4 regions (small fragments < 400 bp were discarded) according to the section “Clean Up Libraries” in the Illumina protocol [54] with minor modifications. Double-size selection was performed for the V4-V5 region (discarded fragments > 670 bp and < 400 bp) following the Ampure XP bead upper & lower cut protocol of DNA Technologies Core. The integrity of the fragments was checked using 1.5% agarose gel and the protocol is provided in the supplementary S3. The Nextera XT Index Kit v2 sets A, B, C (24 indexes, 384 samples; Illumina) were used for dual-indexing. Index PCR protocol and cycling conditions were optimized for the two Zurich datasets and details are provided in the supplementary material S3. The Index PCR-product was purified using the same clean up protocols as described above. The library was quantified using QubitTM dsDNA HS Assay Kit (Invitrogen, Carlsbad, CA, USA) on a Qubit^TM^ Flex Fluorometer following manufacturer’s protocols. The libraries were normalised to 4 nM (diluted with low-TE buffer), pooled and denaturation was conducted following the Illumina 16s metagenomics guide. The final pooled library was diluted to either 6pM, 10 pM or 14 pM and 5% or 10% PhiX control was used depending on the run. Paired-end reads of 2×300 bp or 2x250bp, depending on the run were obtained using Illumina MiSeq FGx Reagent micro kits (600-cycles), MiSeq Reagent Kit V3 (600-cycles) or Illumina MiSeq nano kits (500-cycles) on the Illumina MiSeq sequencing platform (Illumina, Inc., Hayward CA, USA).

### Read data processing for all datasets

Primers were removed for the Dobay et al. (2019) and the V1V3 Zurich dataset 1 using cutadapt (v3.5) and using cutadapt (v4.0) for a subset of V1V3 and V4V5 Zurich dataset 2. For the remaining V3V4 and V4V5 Zurich dataset 2, primers were removed using the trimLeft option in DADA2^42^. All datasets were processed using DADA2 pipeline (version 1.16) with dataset specific quality filtering parameters summarized in the supplementary material S3. Only forward reads were used for V1V3 samples from the Zurich dataset 1 due to the poor quality of reverse reads. ASV abundance tables were clustered into OTUs at 97% similarity threshold against the SILVA database (v138, 2019) for each dataset separately. The SILVA database was also clustered at 97% using RESCRIPt plugin from QIIME2 ^43–46^. Decontam (v1.16) was used for all datasets separately on reagent blanks with the prevalence method and threshold of 0.1^47^. OTU abundance tables from all datasets were agglomerated using OTU IDs with MASS, reshape2, dplyr and data.table libraries in R(v4.2.1). Samples with less than 1000 reads were removed. The phylogenetic tree was constructed using the full length sequences of all the OTU IDs from the single-source control samples (457 samples). Full length sequences for all OTU IDs were extracted from the same 97% clustered SILVA database that was used during closed-reference OTU Clustering. Rooted phylogenetic tree was created using MAFFT and FASSTREE using QIIME2 (v2023.5). Further processing was conducted using the phyloseq package in R(v4.2.1). OTU abundances were normalized using total sum scaling and beta diversity with bray curtis and weighted Unifrac distances were explored using phyloseq, vegan, rbiom and ggplot2 packages in R. All Regression analyses were conducted in R(v4.2.1).

### Machine learning with Random Forest Classifier for all datasets

A total of 457 single-source control samples and 6455 OTUs. Control samples from all 4 regions of the 16S rRNA gene and from saliva, semen, skin-penile, skin from hands, urine and vaginal swab. The vaginal fluid and menstrual blood categories were collectively referred to as “vaginal swab” since they share the same microbial ecosystem. The random forest classifier (RFC) was trained with an 80-20 split using the q2-sample-classifier plugin in QIIME2 (v2023.5) using --p-estimator RandomForestClassifier, --p-optimize-feature-selection and --p-parameter-tuning options. The final RFC was trained on a set of 281 OTUs determined by recursive feature extraction in the --p-optimize-feature-selection option and 5-fold cross-validation was conducted with --p-n-estimators 500. Receiver Operating Curves (ROC), Area Under the Curve (AUC) and confusion matrices were generated while running the q2-sample-classifier command. Precision, recall and F1-scores were calculated using the Metrics and caret packages in R(v4.2.1).

Furthermore, another RFC was trained on the entire dataset of 457 samples and a final set of 337 OTUs using the same hyperparameters and same QIIME2 parameters and options as before. Using this classifier, 50 blind single-source samples and 285 mock samples including both controlled and uncontrolled mock samples were predicted. All boxplots and stacked bar plots were generated using ggplot2, tidyr, tibble, ggrepel and dplyr packages in R(v4.2.1) with the assistance of ChatGPT for troubleshooting.

Informative feature analyses were done by first extracting the taxonomy for all 337 OTUs from the same SILVA database as the closed-reference clustering using the QIIME rescript filter-taxa command. The top 10 OTUs per training class (45 distinct OTUs) were obtained from the output of the sample-classifier heatmap plugin in QIIME2. A heatmap was generated containing only these OTUs.

### Manuscript writing

LLMs were employed to refine sentence structure and readability of text throughout the manuscript.

## Declarations

### Ethics statement

All samples for the Zurich datasets were collected under the project number (2021-11c) according to the CEBES guidelines.

### Consent for publication

Consent forms were obtained for all samples in the Zurich datasets under the project number (2021-11c) according to the CEBES guidelines.

### Author Contributions

MS, RK, and NA conceptualized and designed the study. The Zurich datasets were generated by MS, MG, LS (Schuh L.), LW, FJ, TF, and NA at the Department of Forensic Genetics, with the assistance of CH. The study design for the Zurich underwear sampling was inspired by SL. SA, SL, EH, and LS (L.Snipen) contributed datasets and provided interpretation and analytical assistance. Statistical and bioinformatics analyses, as well as overall study interpretations, were conducted by MS, NB, RK, and NA. The initial manuscript was drafted by MS, RK, and NA, with all authors participating in the manuscript review process.

## Acknowledgements

We express our gratitude to Dr. Janko Tackmann for his invaluable support and insightful suggestions regarding our data analyses and study design. Our heartfelt thanks also go to Dr. Adelgunde Kratzer for her exceptional generosity and support. We are immensely grateful to Dr. Karan Khathuria for his assistance with the machine learning analysis.

Additionally, we extend our sincere thanks to Lina Kim for her invaluable support during the manuscript writing and editing process.

