## Supplementary material S1 for "Standardising a microbiome pipeline for body fluid identification from complex crime scene stains"

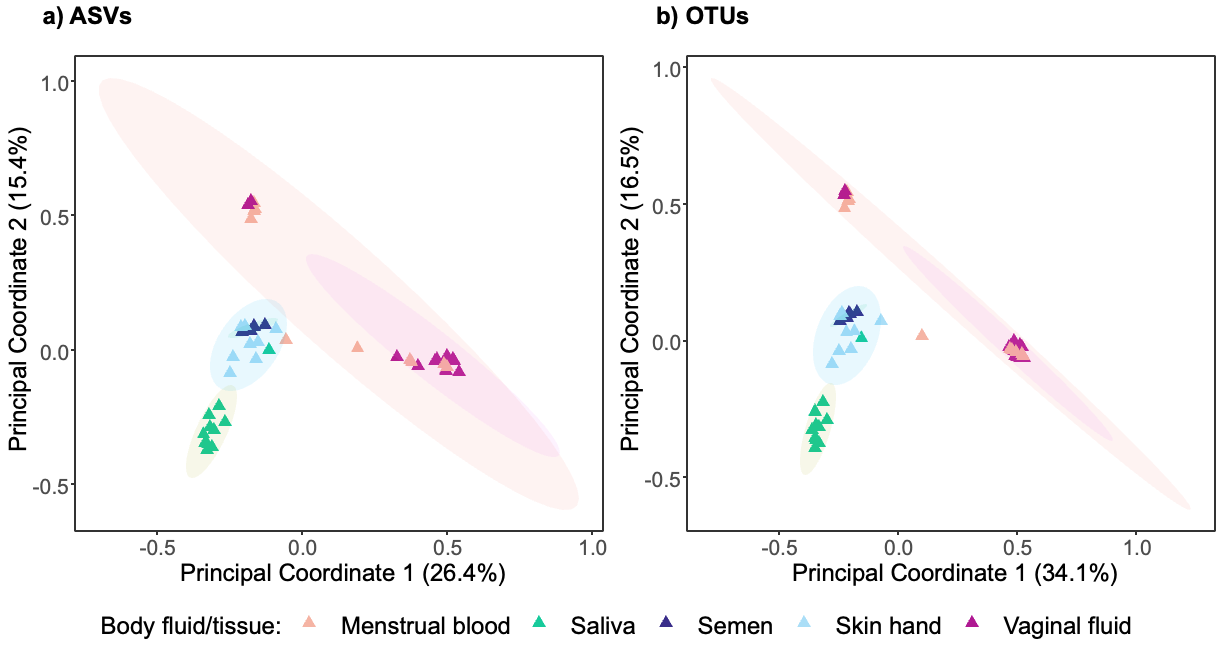

***Figure S1.1 Principal coordinate analysis plots (based on the Bray Curtis distances) showing clustering of body fluid/tissue samples.*** ***a)*** *ASV data for samples from Dobay et al. (n=42, PERMANOVA F_4,41_ = 6.15, r^2^ = 0.38, p = 0.001),* ***b)*** *OTU data clustered at 97% for samples from Dobay et al. (n=42,* PERMANOVA F_4,41_ = 7.64, r^2^ = 0.43, p = 0.001 *).  Body fluid/tissues are colour coded.*

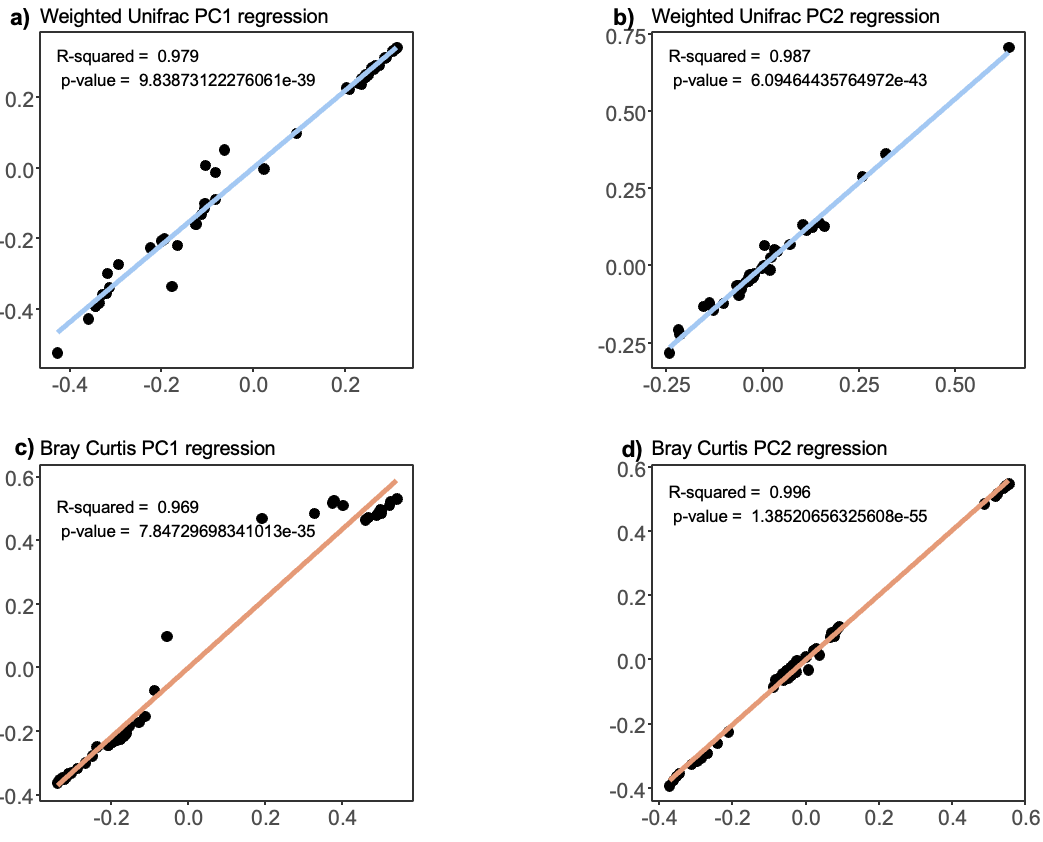

***Figure S1.2 Regression analyses for PC1 and PC2 coordinates for weighted Unifrac and Bray Curtis PCoA plots.*** ***a)*** *Weighted Unifrac PC1 regression,* ***b)****Weighted Unifrac PC2 regression,* ***c)*** *Bray Curtis PC1 regression,* ***d)*** *Bray Curtis PC2 regression.  Regression lines are coloured according to the distances.*

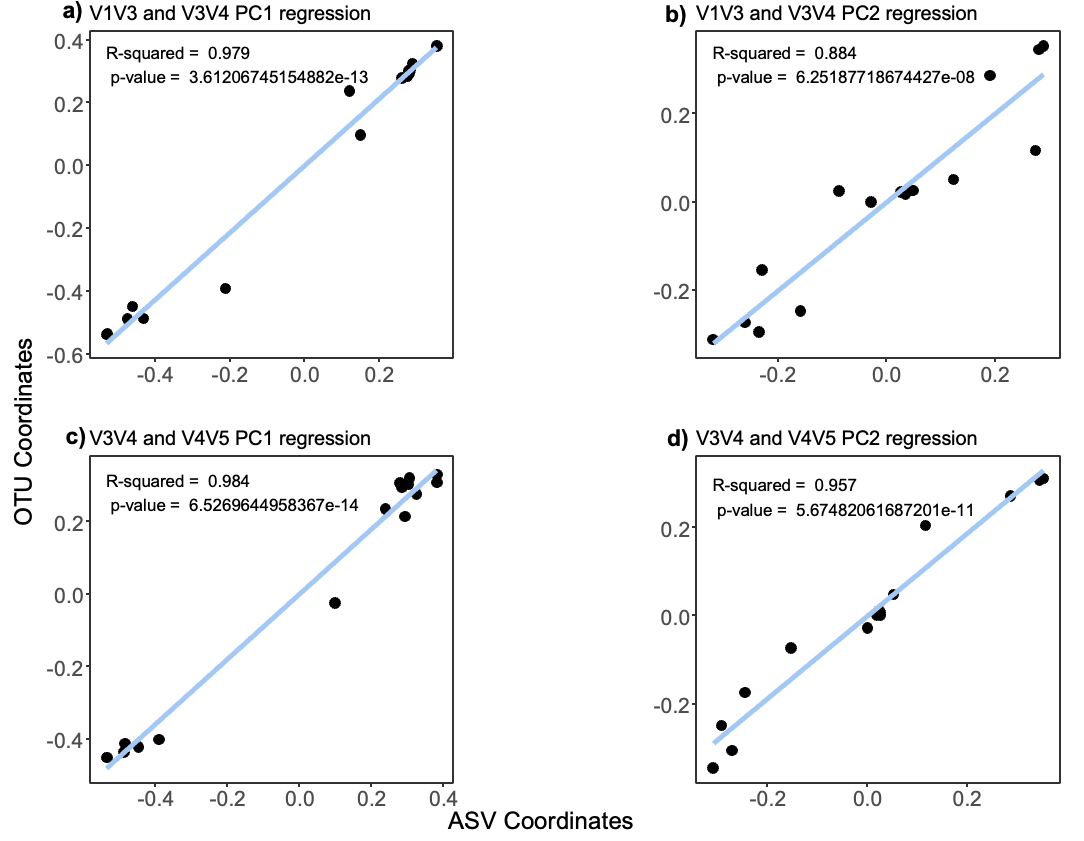

***Figure S1.3 Regression analyses for PC1 and PC2 coordinates for V1V3 and V3V4; V3V4 and V4V5 coordinates.*** ***a)*** *V1V3 and V3V4 PC1 regression,* ***b)****V1V3 and V3V4 PC2 regression,* ***c)*** *V3V4 and V4V5 PC1 regression,* ***d)*** *V3V4 and V4V5 PC2 regression.*

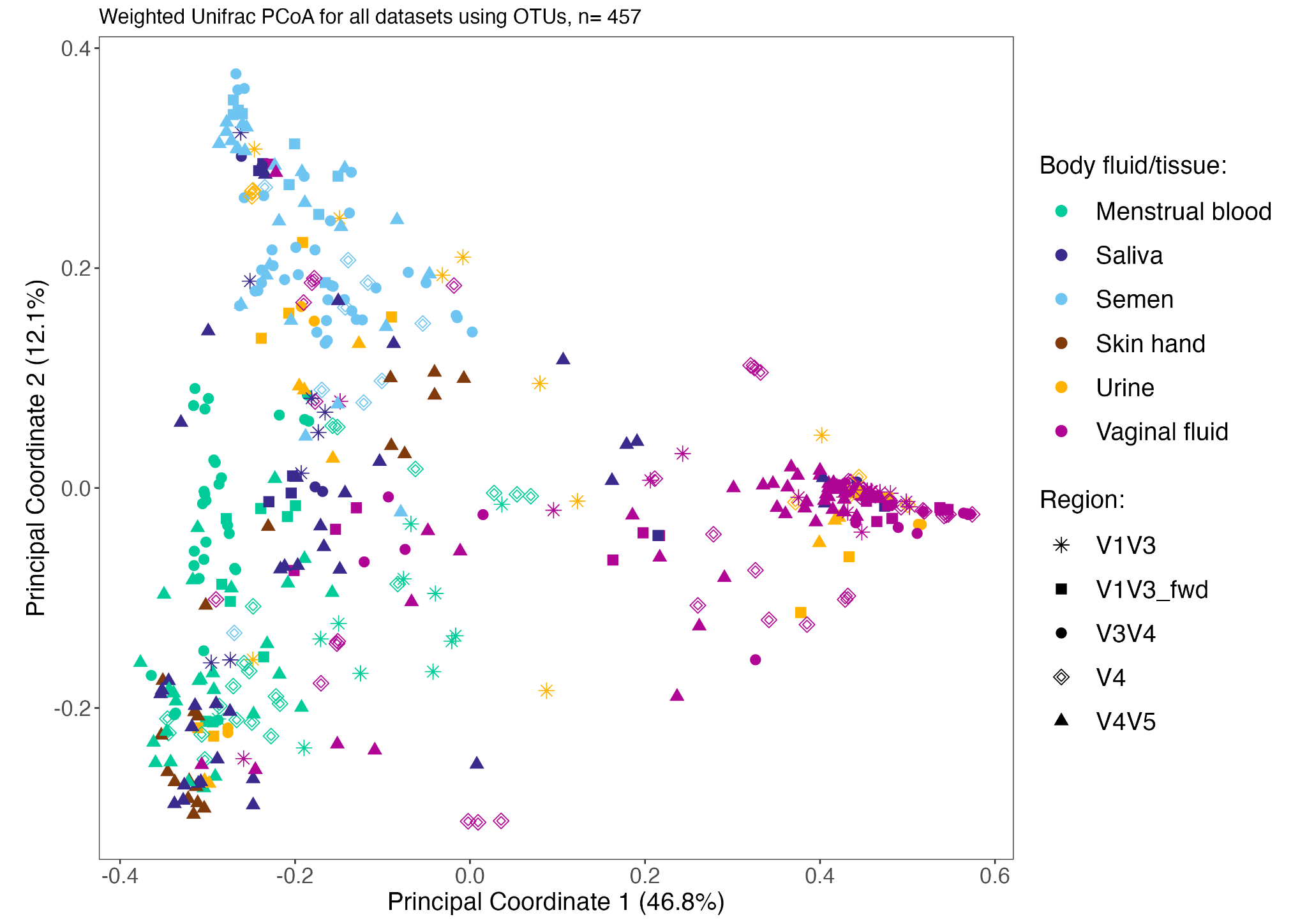

***Figure S1.4 Principal coordinate analysis plots for data from V1V3, V3V4, V4 and V4V5 16S rRNA gene regions using OTUs (97%) and weighted unifrac distances for 457 samples.*** *(PERMANOVA with body-site F_6,450_ = 55.33, r^2^ = 0.42, p = 0.001, PERMANOVA with region F_4,452_ = 6.82, r^2^ = 0.06, p = 0.001)*

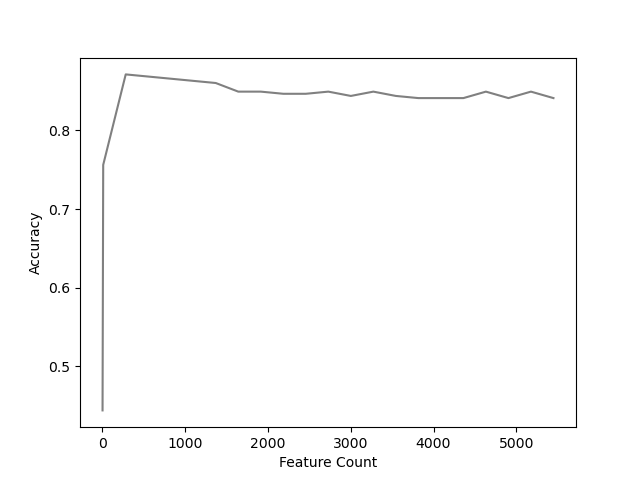

***Figure S1.5 Recursive feature extraction plot for the classifier trained on 365 samples.***

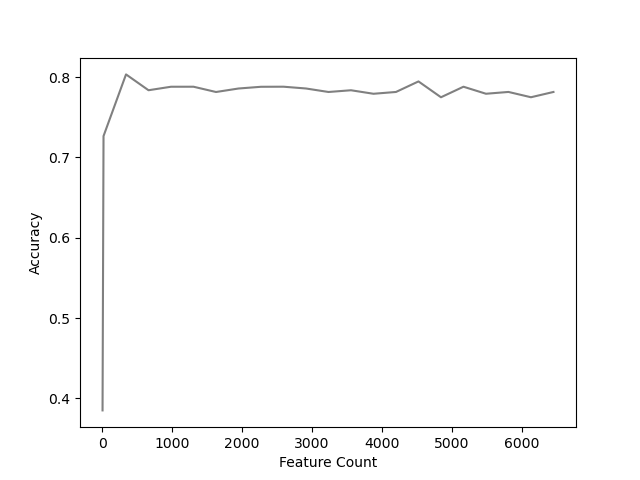

***Figure S1.6 Recursive feature extraction plot for the classifier trained on 457 samples.***

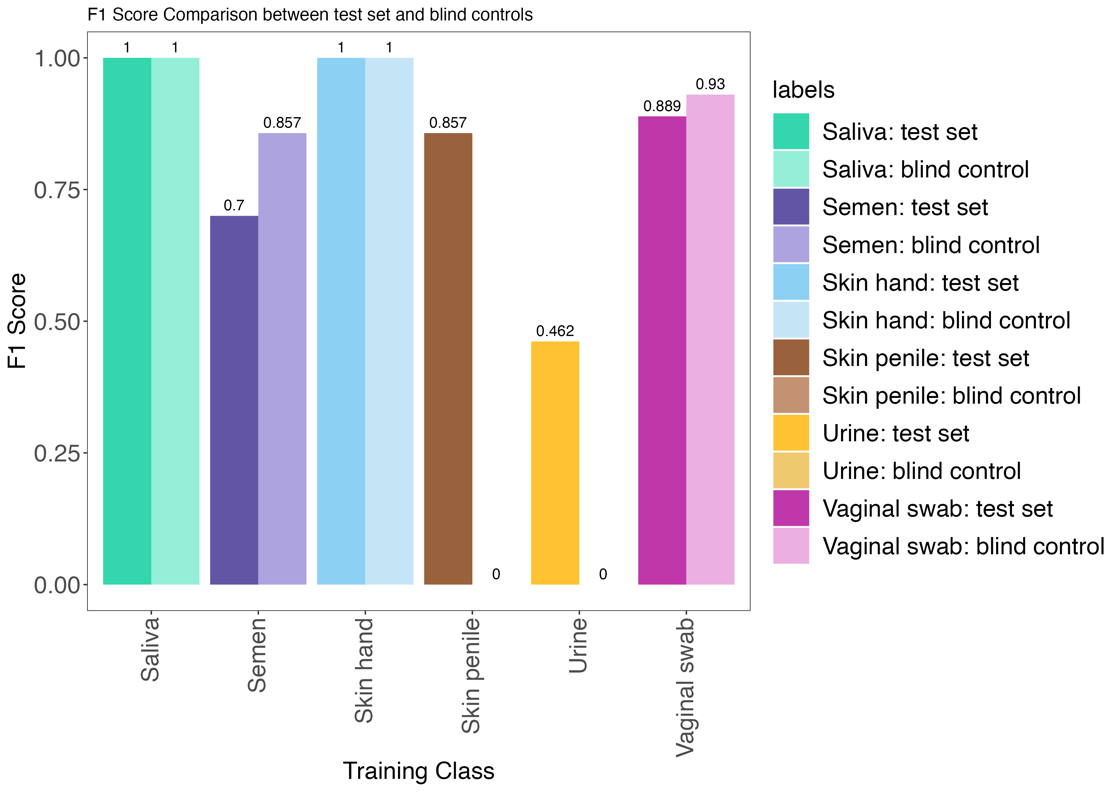

***Figure S1.7 Classifier performance comparison of the classifier trained on 365 samples with test samples vs classifier trained on 457 samples with blind control samples.*** *Barplots depict F1 scores per class and are color coded. Vaginal swab refers to both vaginal fluid and menstrual blood.*

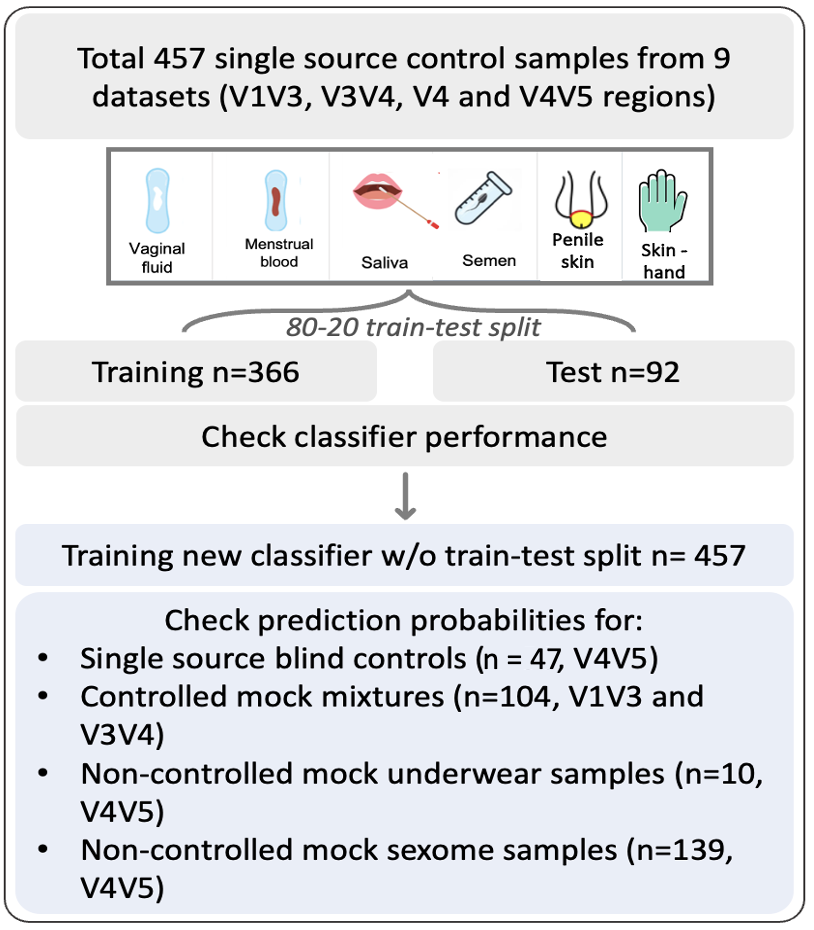

***Figure S1.8 Illustrates an overview of datasets used in classifier training and testing.***
