## Supplementary material S3 for "Standardising a microbiome pipeline for body fluid identification from complex crime scene stains"

Generation of ZIFM/Zurich datasets

| **ZIFM/Zurich dataset1** | **ZIFM/Zurich dataset2** |
| --- | --- |
| **Overview of samples** | |
| Single source control samples from saliva, vaginal fluid, menstrual blood, semen and urine.  Controlled mock mixed-source samples: saliva-semen, vaginal fluid-urine, menstrual blood- semen and vaginal fluid-semen, 9 samples in each category. | Single source control samples from saliva, vaginal fluid, menstrual blood, semen, urine and skin from hand.  Uncontrolled mock samples from underwear:  10 underwear samples were collected from women after wearing the garment for 24h. Participants were required to fill a questionnaire.  Uncontrolled mock samples from the sexually shared microbiome (sexome) study:  Samples from vaginal fluid, semen and penile skin from 22 couples. |
| **Sample collection** | |
| Prionics forensix BasicDry Evidence Collection Tube (Prionics AG, Schlieren Zurich, Switzerland) | Controls:  Sterile cotton swabs (Deltalab, Millian, Nesselnbach, Switzerland)  Uncontrolled mock samples from sexome study:  “ForensiXs” cotton tip (Prionics Lelystad BV, Netherlands) swabs and the cotton-tip swabs by Milian (Deltalab, Millian, Nesselnbach, Switzerland) were used. |
| Controls:  Single source control vaginal fluid and menstrual blood were directly collected with swabs. Single source urine, semen, and saliva were first collected in 50ml sterile collection tubes and 50 μl for each were pipetted onto swabs.  Controlled mock mixed-source samples:  Saliva-semen mixture was prepared by pipetting 25 μl saliva and semen onto the same swab. For mixtures containing vaginal fluid or menstrual blood, vaginal fluid fluid or menstrual blood were collected directly on swabs and 25 uL of urine or semen were pipetted onto the swabs. | Controls:  All samples were collected directly or pipetted onto sterile cotton swabs (Deltalab, Millian, Nesselnbach, Switzerland), dried for at least 2h and stored at -20C. Menstrual blood, skin from hand and vaginal fluid samples were collected directly on swabs. For samples from skin from hands, moist swabs (50 μl ddH2O) were rubbed over the fingertips of the strong hand (one finger per swab). Fresh saliva, semen, and urine were collected in an appropriate container before 50 μl was pipetted onto swabs.  Uncontrolled mock samples from underwear:  The area of interest was selected by visual assessment of the biological stain and the area was cut with a sterile scalpel. A third of the cut out was placed into a 1.5 mL eppendorf tube and stored in a -20° C freezer.  Uncontrolled mock samples from the sexually shared microbiome (sexome) study:  Vaginal fluid and penile skin samples were directly collected on swabs. For penile skin collection, a pre-moistened swab was used to swab along the coronal sulcus of the penis. Semen samples were collected in sterile 50 mL tubes and either 50uL of semen was pipetted onto the swabs or participants were asked to dip the tip of the swab into the collection tubes for 5-10 seconds. All samples were dried for 2 hours and then extracted as soon as possible after collection. |
| **DNA extraction** | |
| QIAamp BiOstic Bacteremia DNA Kit (Qiagen, Germany)  -modified by adding the swab to the PowerBead tubes containing 450 μl Solution MBL and incubating at 10 min at RT and then following the manufacturer’s protocol  DNAeasy PowerSoil Pro Kit (Qiagen, Germany)  -modified by placing the swab in the PowerBead tube containing 800 µl CD1 solution and incubating it for 10 min at RT and then following the manufacturer’s protocol | DNeasy PowerSoil Pro Kit (Qiagen, Venlo, Netherlands)  QIAamp DNA Mini Kit (Qiagen, Venlo, Netherlands)-DNA purification from buccal swabs (Spin protocol) was used both with and without modifications.  -For every swab, the cotton was separated from the wood with a sterile scalpel and placed into a 2 ml tube (Investigator Lyse & Spin Basket Kit, Qiagen, Venlo, Netherlands)  -modified by adding a heating step of 95C for 15 min at 500 rpm in the thermoshaker at the lysis step.  PrepFiler BTA Automated Forensic DNA Extraction Kit (Thermo Fisher Scientific, Waltham, MA, USA). |
| **DNA quantification** | |
| Quantus Fluorometer (Promega, Inc., Madison WI, USA) was used to quantify the DNA concentration of the DNA extracts according to the manufacturer’s protocol. | FemtoTM Bacterial DNA Quantification Kit (Zymo Research, Irvine, CA, USA) or  Protocol described by Seashols-Williams et al. (2018) using the PerfeCTa SYBR Green SuperMix (Quantabio, Beverly, MA, USA) and the ZymoBIOMICS™ Microbial Community DNA Standard D6306 (Zymo Research, Irvine, CA, USA). Based on the Ct numbers, samples were divided into high and low yield. High yield samples were all samples that had Ct numbers =< 17 and 25 cycles were used for these samples. Low yield samples were samples that had a Ct value of >= 17 and 32 cycles were used for these samples together with negative controls. |
| **Library preparation: primers** | |
| V1-V3 primers: F27/R534 (5′-AGAGTTTGATCCTGGCTCAG-3′, 5′- ATTACCGCGGCTGCTGG -3′), Sigma (Darmstadt, Germany).  The V1V3 adapters were modified by including degenerate bases before the primer sequence. | V1V3, V3V4 and V4-V5 regions of the 16S rRNA gene.  V1V3 primers: F27/R534 (5′-AGAGTTTGATYMTGGCTCAG-3′, 5′-ATTACCGCGGCTGCTGG-3′)  The V1V3 adapters were modified by including degenerate bases before the primer sequence and were ordered from Microsynth AG (Balgach, Switzerland).  V3V4 primers: F341/R806 (5′-CCTACGGGNGGCWGCAG-3’,5’-GGACTACHVGGGTWTCTAAT-4’)  V4V5 region primers: F806/R926 (5’-GTGYCAGCMGCCGCGGTAA-3’, 5’-CCGYCAATTYMTTTRAGTTT-3’) |
| **Amplicon PCR** | |
| 25 µL of PCR mixture = 1 µL DNA 12.5uL of the 2X Phusion Hot Start II High-Fidelity PCR Master Mix + 9uL nuclease free water + 1.25uL of the three 10uM forward and reverse primers.  The PCR cycling conditions used were 30 s at 98°C for the initial incubation followed by 10 s at 98°C, annealing 30 s at 57°C , extension 25s at 72°C, a final extension for 10 min at 72°C and hold for at 4°C.  Single source control samples: Menstrual blood, Vaginal fluid, Saliva, negative controls - amplified for 22 cycles  Single source control samples: Urine and Semen- amplified for 32 cycles  Mixtures: amplified for 33 cycles | 25 µL of PCR mixture = 1-5µL DNA, 12.5uL of the KAPA HiFi HotStart ReadyMix PCR Kit (Roche, Basel, Switzerland), 1.25uL of 0.5uM forward and reverse primers and the remaining volume was made up to 25uL with nuclease free water.  The PCR cycling conditions used were 3 min at 95°C for the initial denaturation followed by 20 s at 98°C, annealing 30s at 55-58°C , extension 25s at 72°C, a final extension for 10 min at 72°C and hold for at 4°C.  Depending on the high or low yield samples determined by the qPCR, 25 or 32 cycles were used. |
| **Agarose gel** | |
| 1.5% agarose gel with 5 µL GelRed, 3 µL PCR product, 2.5 µL distilled water and 3 µL DNA loading dye per well and compared against a 100 bp allelic DNA ladder.  The gel was run at 110 V for 40 min. | 1.5% agarose gel with 5 µL GelRed, 3 µL PCR product, 2.5 µL distilled water and 3 µL DNA loading dye per well and compared against a 100 bp allelic DNA ladder.  The gel was run at 110 V for 40 min. |
| **Clean-up 1** | |
| AMPure XP beads (AMPure XP, Beckman Coulter) were used with a ratio of 0.8-fold volume of AMPure XP beads per volume of PCR product. | AMPure XP beads (AMPure XP, Beckman Coulter) were used.  V1-V3 and V3-V4 samples: Single-size selection was performed (small fragments < 400 bp were discarded) according to the section “Clean Up Libraries” in the Illumina protocol with minor modifications.  V4-V5 samples: Double-size selection was performed (discarded fragments > 670 bp and < 400 bp) following the “Ampure XP bead “upper & lower cut” protocol” of DNA Technologies Core. |
| **Index PCR** | |
| The Nextera XT Index Kit (24 indexes, 96 samples; Illumina) was used for indexing. Index PCR reactions were performed in a total volume of 25 μL with 2.5 μL molecular grade water, 12.5 μL of 2X Phusion Hot Start II High-Fidelity PCR Master Mix, 5 μL of PCR product and 2.5 μL of each of indexing primers.  PCR cycling conditions for the index PCR were 30s at 98°C for initial incubation, followed by 12 cycles of denaturation for 10s at 98°C, annealing for 30s at 55°C, extension for 20s at 72°C, a final extension for 5min at 72°C and hold at 4°C. | The Nextera XT Index Kit v2 sets A, B, C (24 indexes, 384 samples; Illumina) were used for indexing. Index PCR reactions were performed in a total volume of 25 μL with 12.5uL of the KAPA HiFi HotStart ReadyMix PCR Kit (Roche, Basel, Switzerland), 7.5 μL of PCR product and 2.5 μL of each of indexing primers.  PCR cycling conditions for the index PCR were 3 min at 98°C for initial denaturation, followed by 12 cycles of denaturation for 20s at 98°C, annealing for 30s at 55°C, extension for 25s at 72°C, a final extension for 1 min at 72°C and hold at 4°C. |
| **Clean-up 2** | |
| AMPure XP beads (AMPure XP, Beckman Coulter) were used with a ratio of 0.6-fold volume of beads per volume of PCR product. | AMPure XP beads (AMPure XP, Beckman Coulter) were used.  V1-V3 and V3-V4 samples: Single-size selection was performed (small fragments < 400 bp were discarded) according to the section “Clean Up Libraries” in the Illumina protocol with minor modifications.  V4-V5 samples: Double-size selection was performed (discarded fragments > 670 bp and < 400 bp) following the “Ampure XP bead “upper & lower cut” protocol” of DNA Technologies Core. |
| **Library concentration** | |
| The library concentration was measured using Spark 10M Multimode Microplate Reader (Tecan) and Qubit dsDNA BR assay (ThemoScientific) using manufacturer’s protocols. | The library was quantified using QubitTM dsDNA HS Assay Kit (Invitrogen, Carlsbad, CA, USA) on a QubitTM Flex Fluorometer following manufacturer’s protocols. |
| **Sequencing** | |
| Normalization and pooling of the libraries were performed with the pipette robot (Liquid Handling Statin, Brand). A pooled library of 3.03 nM was denatured and diluted to 14 pM and a 10% PhiX control (20 pM) was used. Paired-end reads (2×300 bp reads) were obtained using the MiSeq Reagent Kit V3 (600-cycles) on the Illumina MiSeq sequencing platform (Illumina, Inc., Hayward CA, USA). | Normalisation, pooling and denaturation was conducted following the Illumina 16S protocol. The libraries were normalized to 4 nM (diluted with low-TE buffer) and pooled. The final pooled library was diluted to 6pM or 10 pM and 5% or 10% PhiX control was used depending on the run. Paired-end reads (2×300 bp reads) were obtained using Illumina MiSeq FGx Reagent micro kits (600-cycles) on the Illumina MiSeq sequencing platform (Illumina, Inc., Hayward CA, USA). |

**Exact primer sequences**

| **Dataset** | **Fwd primer** | **F_sequence** | **F_pr_len** | **Rev_primer** | **R_sequence** | **R_pr_len** | **Region** |
| --- | --- | --- | --- | --- | --- | --- | --- |
| ZIFM Dataset1 | 27F | AGAGTTTGATCCTGGCTCAG | 20 | 534R | ATTACCGCGGCTGCTGG | 17 | V1V3 |
| ZIFM Dataset2 | 27F | AGAGTTTGATYMTGGCTCAG | 20 | 534R | ATTACCGCGGCTGCTGG | 17 | V1V3 |
| Hanssen et al. (2017) | PRK341F | CCTAYGGGRBGCAACAG | 17 | PRK806R | GGACTACNNGGGTATCTAAT | 20 | V3V4 |
| ZIFM Dataset2 | 341F | CCTACGGGNGGCWGCAG | 17 | 806R | GGACTACHVGGGTWTCTAAT | 20 | V3V4 |
| Ahannach et al. | 515F | GTGCCAGCMGCCGCGGTAA | 19 | 806R | GGACTACHVGGGTWTCTAAT | 20 | V4 |
| Dobay et al. (2019) | 518F | CCAGCAGCYGCGGTAAN | 17 | 926R1 | CCGTCAATTCNTTTRAGT | 18 | V4V5 |
|  |  |  | 17 | 926R3 | CCGTCAATTTCTTTGAGT | 18 |  |
|  |  |  | 17 | 926R4 | CCGTCTATTCCTTTGANT | 18 |  |
| Williams et al. | 515F | GTGYCAGCMGCCGCGGTAA | 19 | 806R | GGACTACNVGGGTWTCTAAT | 20 | V4 |
| Meisel et al. | 515F | GTGCCAGCMGCCGCGGTAA | 19 | 806R | GGACTACHVGGGTWTCTAAT | 20 | V4 |
| ZIFM Dataset2 | 515F | GTGYCAGCMGCCGCGGTAA | 19 | 926R | CCGYCAATTYMTTTRAGTTT | 20 | V4V5 |

**DADA2 parameters for all datasets**

ALL datasets: maxN=0, maxEE=c(2,4), truncQ=2

ZIFM_dataset 1: V1-V3 (Larissa_Schuh): trunclen = c(270,260)

ZIFM_dataset 1: V1-V3 (Larissa_Schuh run2_mix): trimRight = c(10,20)

ZIFM_dataset 2: V3-V4 (Larissa Walser): picked trim left= c(17,20) and trim right= c(0,15)

ZIFM_dataset 2: V4-V5 (Larissa Walser): truclen = c(270,180)

ZIFM_dataset 2: V4-V5 (Larissa Walser TsD) Run1 & Run2: trim left = c(19,20) and trim right =c(20,100)

ZIFM_dataset 2: V4-V5 (Tamara Flury): trim left = c(19,20) and trunclen 220,170

ZIFM_dataset 2: V4-V5 (Fardin Javanmard): trimLeft = c(19,20) and trimRight = c(20,100)

V3-V4 (Hanssen et al.): trunclen = c(260,220)

V4 (Lebeer et al.): trunclen = c(220, 210)

V4 (Meisel et al.): no trunclen no trim

V4 (Williams et al.): truncLen = c(220,220)

V4-V5 (Dobay et al.): trunclen = c(220,220)
